# Non-canonical apical constriction shapes emergent matrices in *C. elegans*

**DOI:** 10.1101/189951

**Authors:** Sophie S. Katz, Chloe Maybrun, Hannah M. Maul-Newby, Alison R. Frand

## Abstract

Specialized epithelia produce apical matrices with distinctive topographies by enigmatic mechanisms. Here, we describe a holistic mechanism that integrates cortical actomyosin dynamics with apical matrix remodeling to pattern *C. elegans* cuticles. Therein, axial AFBs appear near the surface of lateral epidermal syncytia during an interval of transverse apical constriction (AC). AC gives rise to three temporary semi-circular cellular protrusions beneath a provisional matrix (sheath). In turn, sheath components pattern durable ridges along the midline of adult cuticles (alae). We propose that forces generated by AC are relayed via the sheath to sculpt the the acellular adult cuticle manifest several hours later. Furthermore, we provide evidence that circumferential actin filament bundles (CFBs) near the surface of the adjacent syncytia (hyp7) are largely dispensable for the propagation of annular cuticle structures from one larval stage to the next. Rather, the temporary CFBs extend from actin bundles overlying body wall muscles, which are situated between *Ce*. hemidesmosomes. Similar mechanisms may contribute to the morphogenesis of integumentary organs in higher metazoans.

## INTRODUCTION

Polarized epithelial cells give rise to myriad architecturally distinct organs and appendages with specialized functions. These structures either include or adjoin apical extracellular matrices (aECMs) in contact with the environment. Examples in humans include stereocilia on auditory hair cells and microvilli on kidney and intestinal epithelia, which are projections from live cells embedded in matrices; hair, which is an amalgam of keratinized cells and extracellular molecules; and tooth enamel, which is an acellular, mineralized matrix. Mutations that affect related matrix proteins cause deafness, polycystic kidney disease and abnormal teeth (Devuyst et al., 2017; Legan et al., 2005; Masuya et al., 2005; Muller and Barr-Gillespie, 2015). Moreover, carcinomas co-opt physiologic mechanisms that regulate the turnover of apical matrices (Braidotti et al., 2004; Jonckheere et al., 2010). Despite the medical significance of the integument, the cell and molecular mechanisms that sculpt apical matrices are not well understood.

Development of many epithelial tissues involves selective constriction of the apical surface of one or more cells (Heer and Martin, 2017; Iruela-Arispe and Beitel, 2013). Resulting changes in cell shape promote morphogenetic processes ranging from gastrulation to formation of bronchial buds and branches (Kim et al., 2013; Martin et al., 2010; Vasquez et al., 2014). The mechanical networks that drive apical constriction (AC) include actomyosin filaments and/or bundles that generate contractile forces; cell-cell junctions that link actomyosin filaments in adjacent cells; and cortical flows, which promote assembly of actin filaments at regions under tension (Hannezo et al., 2015). Fine spatial and temporal regulation of actomyosin contractility achieves the correct, contextual shape changes. Although these mechanical networks are well characterized (Martin and Goldstein, 2014), the extent to which they shape apical matrices is unclear. Conversely, the contributions of apical matrices to tissue-level outcomes of AC remain enigmatic. This gap reflects our relatively limited knowledge about biochemical and biophysical properties of apical ECMs, as compared with basement membranes.

The key cell and molecular components driving apical constriction are conserved between nematodes and mammals. As depicted in Figure 1, the epidermis of *C. elegans* consists mainly of the lateral stem cell-like seam cells and the adjacent hyp7 syncytium. The seam cells undergo asymmetric division early in each larval stage. The posterior daughters remain in the seam while the anterior daughters fuse with hyp7. During the larval-to-adult transition, seam cells undergo homotypic fusion, becoming bilateral syncytia, and enter persistent cell-cycle quiescence (Sulston and Horvitz, 1977). By then, the hyp7 syncytium includes 139 nuclei (Yochem et al., 1998). Hyp7 has four discernable sub-regions: thick bilateral regions; dorsal and ventral ridges; and thin segments anchored to underlying body wall muscles by trans-epidermal attachments called fibrous organelles (Bosher et al., 2003). Together, the seam and hyp7 synthesize most of the body cuticle.

**Figure 1.**
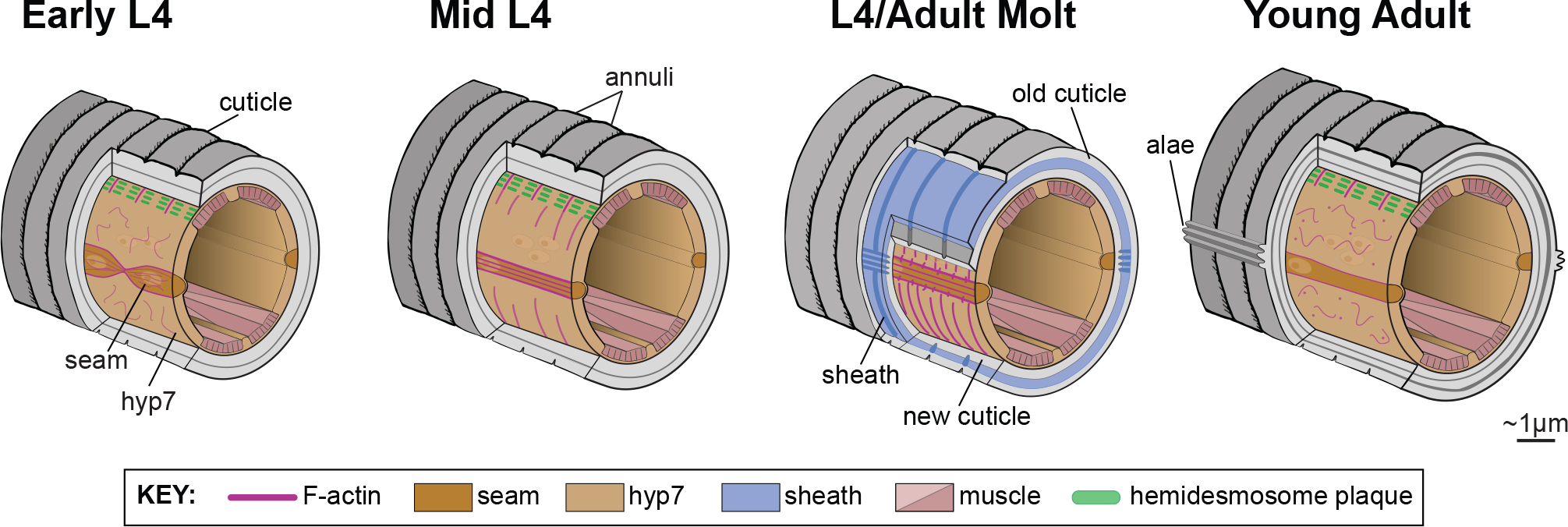
Revised model for progression of the L4/A molt and formation of the alae. Illustrations of *C. elegans* body sections depict changes in the epidermis, cortical actin networks therein, and overlying matrices. These drawings encapsulate both existing knowledge and new findings described in this report.

The apical-lateral junctions among epithelial cells of *C. elegans* are analogous to those found in vertebrates and insects (Pasti and Labouesse, 2014). Homologs of α-catenin, ß-catenin, p120-catenin, and e-cadherin, all of which are components of mammalian adherens junctions (AJs), form a complex subjacent to the apical membrane (Costa et al., 1998; Loveless and Hardin, 2012). The worm homolog of the *Drosophila* septate junction component Discs-large and the nematode-specific protein AJM-1 comprise a distinct basal complex (Koppen et al., 2001; McMahon et al., 2001).

The *C. elegans* cuticle is an acellular, multi-layered extracellular matrix mainly composed of collagens (Page and Johnstone, 2007), rather than the polysaccharide chitin found in insects (Fabritius et al., 2011). The first cuticle is produced late in embryogenesis beneath a provisional matrix called the embryonic sheath (Priess and Hirsh, 1986). Each larval-stage cuticle is shed and replaced with a larger structure during the subsequent four molts (Knight et al., 2002; Sulston and Horvitz, 1977). The adult-stage cuticle lasts for weeks, rather than hours, as adults no longer molt.

The prevailing model for the process of molting includes three sequential steps: 1) detachment of the epidermis from the preexisting cuticle (apolysis); 2) synthesis of the new cuticle directly beneath the old; and 3) escape from the old cuticle (ecdysis) (Jenkin and Hinton, 1966; Locke and Huie, 1979; Truman, 1992). Because this model does not invoke a provisional enclosure, it cannot explain how squishy nematodes remain intact, rather than implode. We recently learned that *C. elegans* larvae produce an interim aECM (sheath) between the epidermis and the old cuticle as they begin to molt (Fig. 1). Notably, specific components of the sheath are patterned like the upcoming, rather than passing, cuticle. The sheath and old cuticle are shed together at ecdysis ((Katz et al., 2015), with permission).

Three longitudinal ridges (alae) overlying the syncytial seam are prominent features of the adult-stage cuticle. Because the alae arise above the syncytial seam during the larval-to-adult transition, numerous studies vis-à-vis the succession of temporal cell fates have scored alae formation as a proxy for maturation of the epidermis (Ambros and Horvitz, 1984). Similarly positioned but morphologically distinct alae are found on the cuticles of L1s and dauers, which are stress-induced variants of L3s (Hu, 2007); but not on the cuticles of L2s, L3s or L4s. To our knowledge, the role of actomyosin-based mechanical networks in patterning the alae has not previously been examined. Indeed, the cortical actin networks of larval seam cells and syncytia have not been described.

Scores of evenly-spaced circumferential bands (annuli) separated by narrow furrows are also found in the outermost layers of all 5 stage-specific cuticles, overlying hyp7 and other epithelia (Page and Johnstone, 2007). The spacing of the annuli is thought to be similar in larvae and young adults (Cox et al., 1981b). More recent studies of adult animals carrying mutations that affect body morphology have shown that the frequency of annuli along the long axis of the body scales with size (Essmann et al., 2017). Parallel rows of circumferentially-oriented cortical actin filament bundles (CFBs) assemble and disassemble in the hypodermis while embryos elongate and again while larvae molt. The CFBs of embryos are thought to pattern the annuli of L1-stage cuticles (Costa et al., 1997; Priess and Hirsh, 1986). However, the mechanism that subsequently relays this annular pattern from effete to emergent cuticles is not known.

Here, we describe a novel and potentially conserved morphogenetic mechanism that integrates cortical actin dynamics with apical matrix remodeling. Therein, a pulse of anisotropic apical constriction generates temporary protrusions from epithelia coupled to the provisional sheath, which then patterns durable ridges (alae) on *C. elegans* cuticles. In this non-canonical system, interim matrices serve as key components of mechanical networks that shape integumentary appendages. In addition, we provide evidence that transient CFBs present in the hypodermis of molting animals appear to be elaborations of continual but previously unrecognized cortical actin bundles overlying body wall muscles, rather than form factors for the annuli.

## RESULTS

### Actin and NMYII pattern longitudinal, rather than annular, folds in adult cuticles

To evaluate the role of actin filaments and/or bundles in patterning emergent cuticles, we used both bacterial-mediated RNA-interference (RNAi) and conditional mutations to knockdown actin and non-muscle myosin II (NMYII) while larvae developed and then examined the lateral surface of young adults. Longitudinal ridges overlying seam syncytia (alae) were directly observed by DIC microscopy. Raised circumferential bands overlying hyp7 (annuli) were detected by staining animals with the fluorescent lipophilic dye DiI prior to observation (Schultz and Gumienny, 2012).

Five genes of *C. elegans* encode actin. Epidermal cells and syncytia express *act-1, -2*, *-3* and possibly *-4* but evidently not *act-5* (MacQueen et al., 2005; Sarov et al., 2012; Willis et al., 2006). To simultaneously knockdown *act-1*, *-2*, *-3* and *-4*, we selected a dsRNA trigger complementary to all four transcripts. Further, we customized and applied an established experimental paradigm to selectively knockdown *actin* in the seam or hyp7. This system entails the tissue-specific expression of wild-type *rde-1*, which encodes the worm homolog of Argonaute, in *rde-1(null)* mutants otherwise insensitive to siRNAs (Qadota et al., 2007; Steiner et al., 2009). Epidermis-specific knockdown of *actin* minimized the larval lethality associated with systemic RNAi of *actin* over the 42-hour course of larval development. Attenuated exposure to *actin* dsRNAs for only 30 hours starting in the L2 stage also bypassed this lethality, allowing otherwise wild-type, *actin(RNAi)* larvae to develop into relatively small adults.

Preferential knockdown of *actin* in either the seam or hyp7 led to the formation of grossly misshapen patches of cuticle, which interrupted the three ridges characteristic of adult-stage alae (Fig. 2A). Patches with many tortuous, short ridges and discontinuous longitudinal ridges were observed on 93% of seam-specific *actin(RNAi)* animals and 89% of hyp7-specific *actin(RNAi)* animals (n≥100). Hereafter, we refer to such patches as “mazes.” The most extensive mazes lacked detectable dorsal and ventral ridges. Such deformities were not found on the lateral surface of *rde-l(null)* mutants fed the same bacteria transformed with empty vector. However, mazes were observed on 89% of wild-type animals following attenuated exposure to *actin* dsRNAs (Fig. 2A-B). Some mazes appeared next to large (>5μm) stretches of cuticle devoid of longitudinal ridges. These stretches uniformly aligned with gaps in the subjacent seam syncytium detected by DIC and fluorescence microscopy. These gaps may be attributable to the loss of seam nuclei during aberrant, larval-stage cell divisions. Accordingly, in our scoring rubric mazes were prioritized *post-hoc*, over large gaps and minor deformities found along the midline of any particular worm.

**Figure 2.**
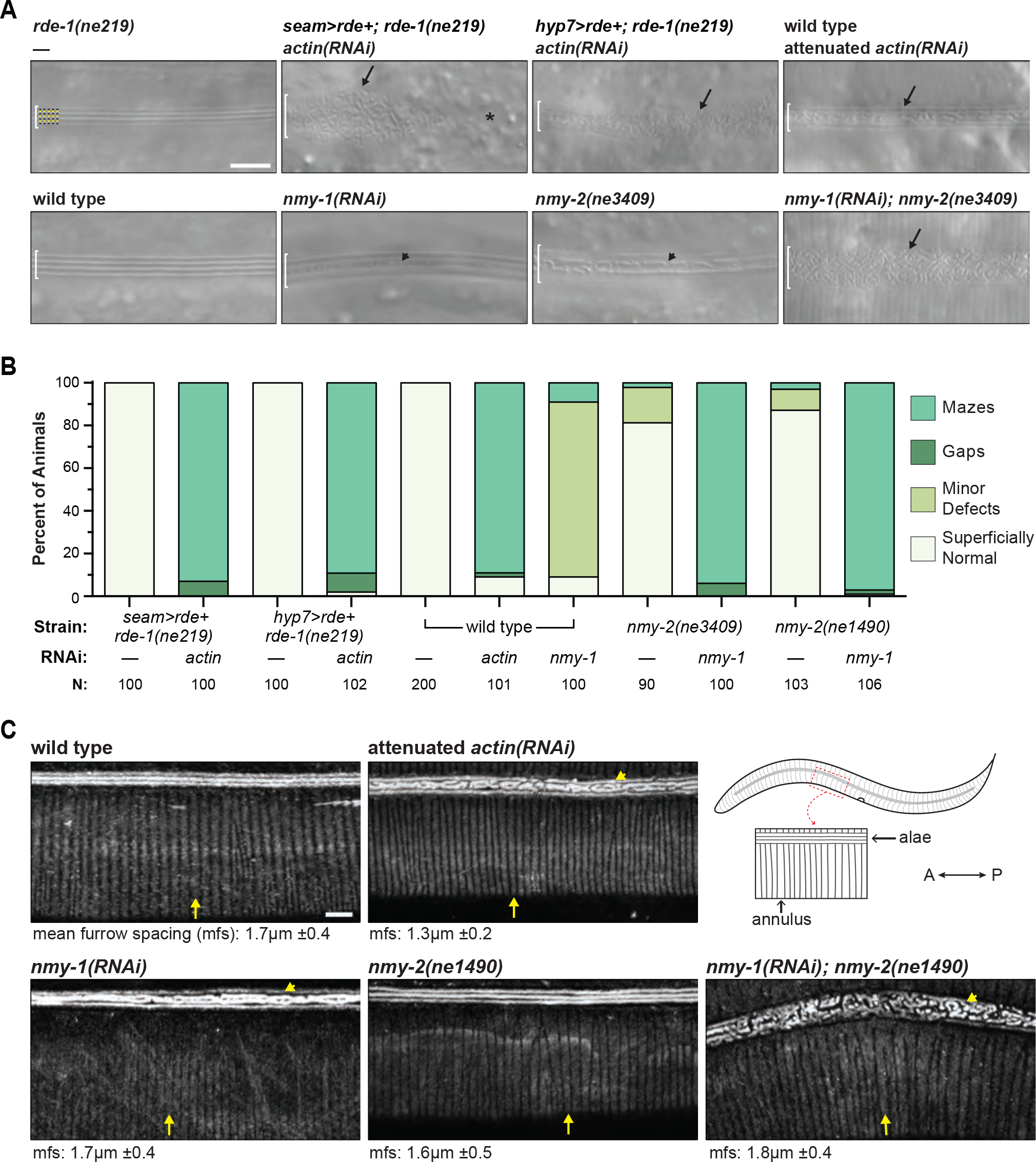
Knockdown of epidermal actin or NMYII results in maze-like deformities on adult cuticles. **A)** Representative DIC micrographs show the lateral surfaces of young adults of the indicated genotypes. The “>” symbols refer to promoter fusions, as described in the methods and Table S1. Brackets demarcate regions above seam syncytia. Yellow and black dashes label presumptive longitudinal ridges (alae) and adjacent valleys, respectively. Arrows and arrowheads point to mazes and minor deformities in alae. Asterisk labels a gap. **B)** Quantitation of alae defects. Values are weighted averages from 2 independent trials. P<0.001 for all pairwise comparisons between control and *actin(RNAi)* or *nmy-1(RNAi)* cohorts; Fisher’s exact test with Bonferroni’s correction for multiple comparisons. **C)** Confocal fluorescence micrographs show Dil-stained cuticles of young adults. Arrows to a single annulus and similar structures; arrowheads point to malformed alae. The mean furrow spacing (±SD) for each genotype is listed underneath corresponding images. Schematic depicts the normal pattern of Dil-staining. Scale bars = 5μm. Strains used were ARF330, ARF408, N2, WM179 and WM180.

Two genes of *C. elegans* encode NMYII heavy chains approximately 47% identical to human NMMHC-IIB, and active in the epidermis (Piekny et al., 2003). We used *nmy-1(RNAi)* and conditional *nmy-2(ts)* alleles to partially inactivate NMYII throughout this study. Mazes, by and large extensive, were observed on 94% of *nmy-1(RNAi); nmy-2(ne3409ts)* double mutants and 97% of *nmy-1(RNAi); nmy-2(ne1490ts)* double mutants raised at restrictive temperature (Fig. 2A-B). Minor deformities in the alae such as small divots were observed on 82% of *nmy-1(RNAi)* single knockdowns and a minority of *nmy-2(ne3409ts)* and *nmy-2(ne1490ts)* single mutants. Collectively, these findings suggest that actomyosin networks within the seam and hyp7 syncytia cooperatively shape the adult-stage alae. The *nmy-1* and -2 genes may act redundantly or cooperatively in this context.

As described, Dil labels raised segments of the cuticle including alae and annuli but does not impregnate adjacent valleys and furrows (Schultz and Gumienny, 2012). As expected, Dil stained scores of evenly-spaced circumferential annuli on dorsolateral and ventrolateral surfaces of wild-type adults (Fig. 2C). Dil also labeled evenly-spaced, annular bands on the surface of *actin(RNAi)* adults, while labeling mazes on the lateral midline. The average spacing of furrows *actin(RNAi)* animals was 1.3μm±0.2 (mean and SD), compared to 1.7μm±0.4 in age-matched controls. This was consistent with the small size of *actin(RNAi)* adults and the reported scaling of these two morphometric features (Essmann et al., 2017). Dil likewise stained evenly-spaced, annular bands on the surfaces of *nmy-1(RNAi)* single knockdowns, *nmy-2(ts)* single mutants, and *nmy-l(RNAi) nmy-2(ts)* double mutants, which had grossly misshapen alae (Fig. 2C). Because Dil consistently saturated the alae, signal in adjacent regions of the cuticle often appeared less intense and more variable among isogenic worms. The formal possibility that these knockdowns lead to subtle defects in the curvature or texture of annuli cannot be eliminated using this approach. Nonetheless, we found no evidence that regenerating the pattern of annuli in adult cuticles requires actin or NMYII, whereas *de novo* patterning of the alae clearly required the corresponding genes. This particular finding contradicts the widely-accepted model that transient CFBs present in hyp7 during molts pattern the annuli of nascent cuticles (Costa et al., 1997).

### AJ components that interact with actin networks shape the adult-stage alae

To evaluate the role of cell-cell junctions in patterning emergent cuticles, we similarly used RNAi and a hypomorphic allele to knockdown key AJ components while larvae developed and then examined the lateral surface of young adults. As described, HMP-1/a-catenin is the actin-binding component of cadherin-catenin complexes (CCCs) that mechanically link the various epidermal cells of *C. elegans* (Costa et al., 1998; Kang et al., 2017; Loveless and Hardin, 2012). The Zonula Occludens (ZO) homolog ZOO-1 cooperatively recruits actin bundles to AJs (Lockwood et al., 2008). We used the substitution mutation *hmp-1(fe4)* because the corresponding protein has decreased affinity for actin but approximately half of the homozygous mutants complete embryonic and larval development (Maiden et al., 2013; Pettitt et al., 2003). We used RNAi of *zoo-1* to sensitize the genetic background. Mazes were observed on the surface of 46% (n=107) of *zoo-1(RNAi); hmp-1(fe4)* double mutants (Fig. 3A-B). Continuous dorsal and ventral ridges framed the majority of these mazes, as observed upon attenuated RNAi of *actin* in wild-type adults (Fig. 2A). Further reduction of *hmp-1* activity may well result in extensive mazes. Dil labeled periodic, annular bands on the dorsolateral and ventrolateral surfaces of all (n=17) *zoo-1(RNAi; hmp-1(fe4)* double mutants, despite the high penetrance of misshapen alae (Fig. 3C). Furrow spacing in the double mutants was 1.4μm±0.4 compared to 1.7μm±0.4 in age-matched wild type controls, consistent with the smaller size characteristic of *hmp-1(fe4)* mutants. The Dil signal and background fluorescence appeared more intense on *hmp-1(fe4)* mutants than wild type adults. This difference may result from relatively inefficient removal of unbound dye through free movement on NGM plates, as the locomotion of *hmp-1(fe4)* adults was visibly impaired. Thus, genetic manipulations that compromise the ability of AJs to transmit forces between the seam and hyp7 abrogated the pattern of adult-stage alae but had little, if any, effect on propagation of the pattern of annuli from L1- to adult-stage cuticles.

**Figure 3.**
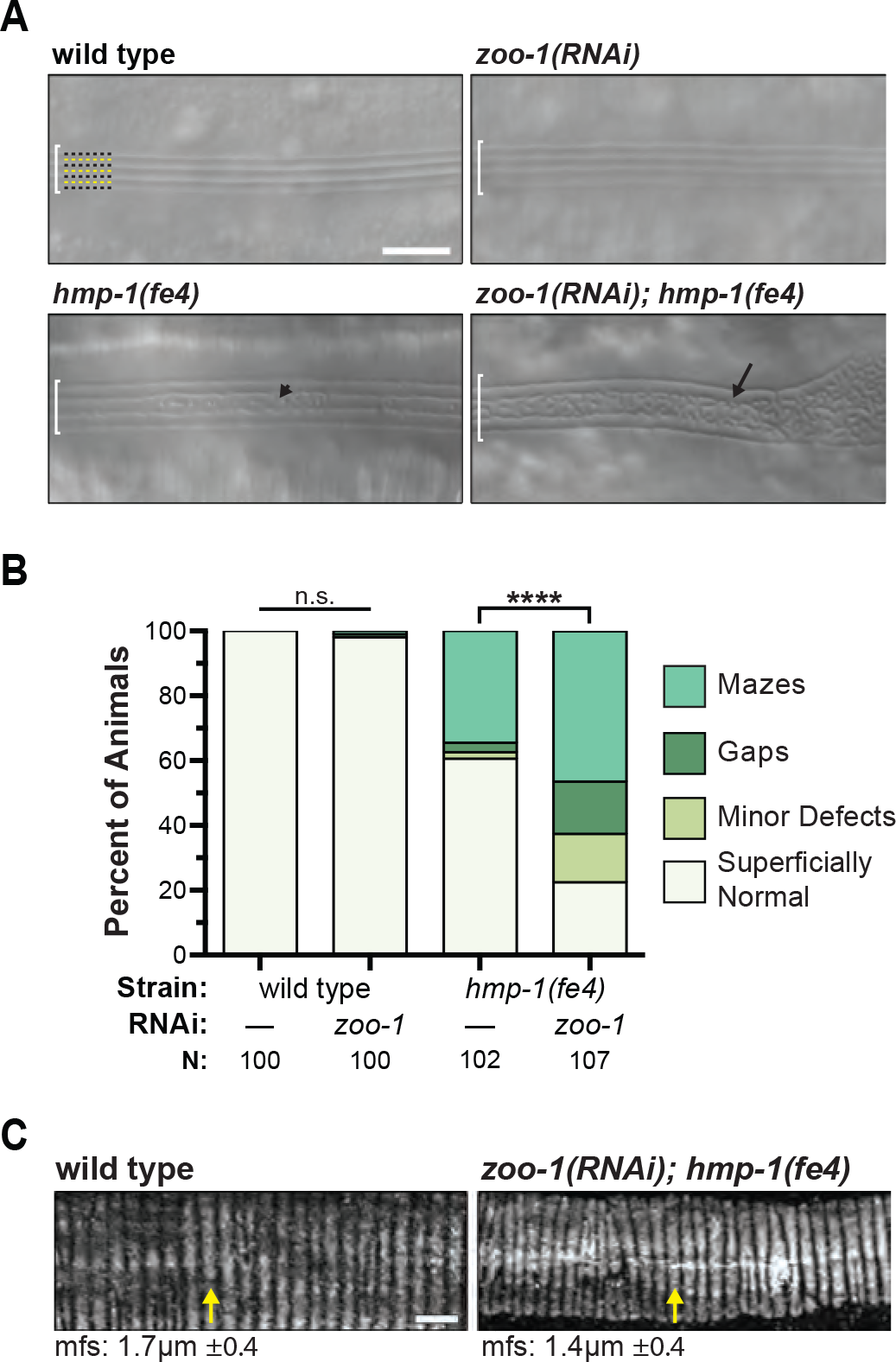
Knockdown of α-catenin and ZO-1 results in mazes on adult cuticles. **A)** As described for Fig. 2, DIC micrographs show segments of lateral cuticles. **B)** Quantitation of alae defects. Values are weighted averages 2 two independent trials. ****P<0.0002, ^n.s.^P>0.99, Fisher’s exact test with Bonferroni’s correction for multiple comparisons. **C)** Confocal fluorescence micrographs show Dil-labeled dorsal-or ventral-lateral surfaces of adult cuticles. Arrows point to an annulus and a superficially similar band. Mfs= mean furrow spacing of corresponding genotype, αSD. Scale bars = 5μm. Strains N2 and PE97 were imaged.

### Cortical actin dynamics linked to pulsatile apical constriction of seam syncytia

To further investigate the role of actin networks and AJs in shaping emergent alae, we tracked the distribution of F-actin at the cortex of seam cells and syncytia in animals developing from the L3 to the adult stage (Fig. S1, Fig. 4). For this purpose, we constructed a genetically-encoded sensor for F-actin comprising the Calponin homology domain (CH) of human Utrophin (UTRN) tagged with GFP and driven by the seam-specific promoter of *elt-5* (Koh and Rothman, 2001). This approach enabled highly-sensitive live-imaging because UTRNCH::GFP binds F-actin selectively and reversibly; in addition, UTRNCH::GFP has no appreciable effect on actin rearrangements when expressed at practical levels (Burkel et al., 2007; Moores and Kendrick-Jones, 2000). To capture both cell and molecular dynamics germane to morphogenesis of the alae, we combined AJM-1::mCHERRY, an established marker for adherens junctions (Koppen et al., 2001), with the seam>UTRNCH::GFP sensor. To achieve fine temporal resolution, we isolated precisely staged transgenic nematodes as follows: We selected ∼60 individuals transiting the L3/L4 molt from synchronously developing populations and watched them; the time when a given worm emerged in L4 was set to zero; we then collected cohorts of ∼7 worms at regular ∼1hr intervals and imaged both GFP and mCherry near the apical surface of the lateral epidermis using confocal fluorescence microscopy.

**Figure 4.**
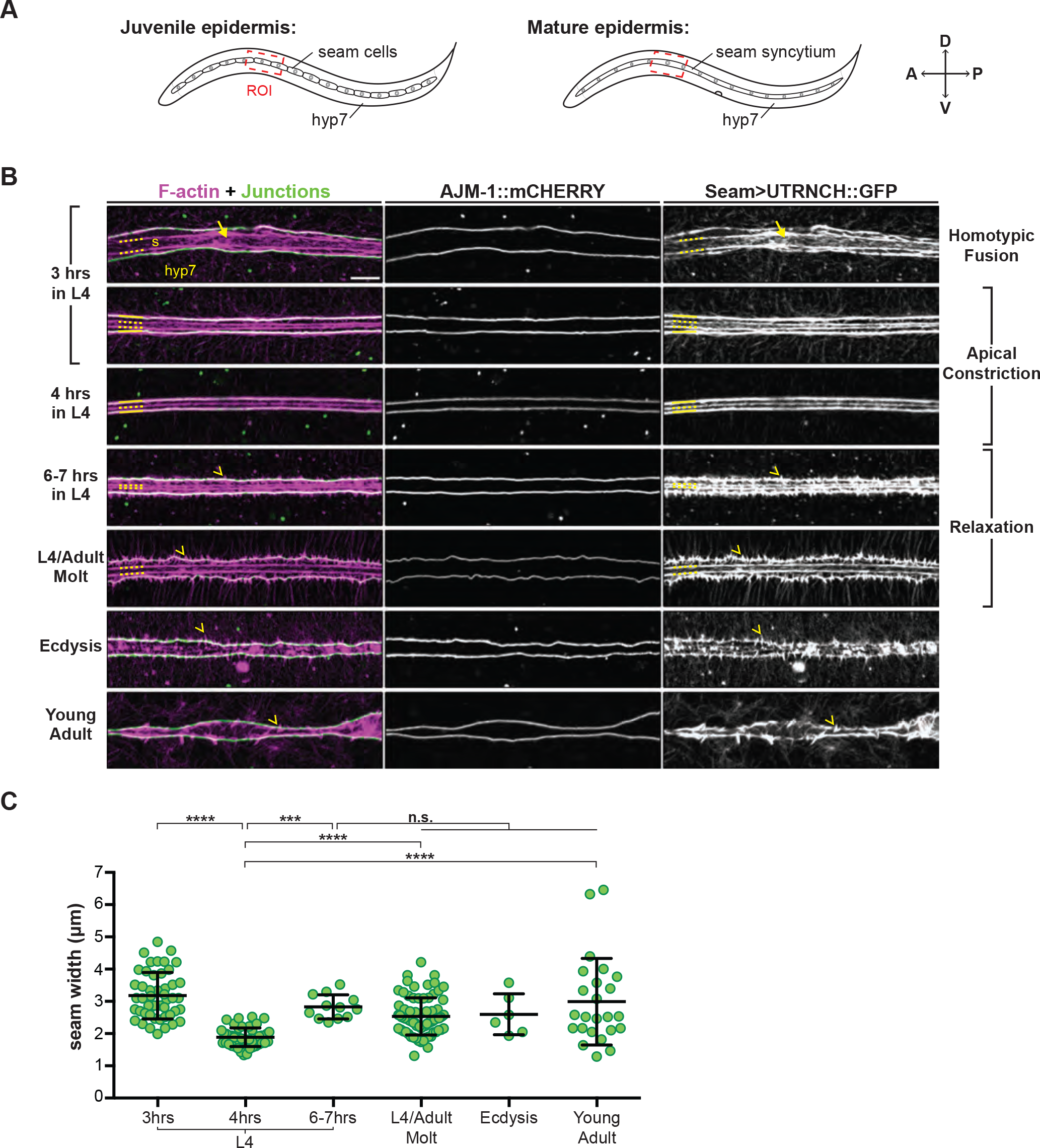
Actin dynamics linked to pulsatile constriction of the seam cortex. **A)** Diagrams depict the seam cells and syncytia of juvenile and mature *C. elegans*. Images in B and C generally correspond to the indicated region of interest (ROI). **B)** Representative confocal fluorescence projections show signals from seam>UTRNCH::GFP (false colored magenta) and/or the AJ marker AJM-1::mCHERRY (false colored green). Yellow lines label longitudinal actin bundles at the dorsal and ventral margins. Dashes label medial-axial actin bundles within seam syncytia (s). Arrow points to cortical actin mesh. Chevrons point to F-actin spikes orthogonal to seam/hyp7 margins. **C)** Quantitation of seam width measured as distance between AJM-1::mCHERRY-marked junctions. Lines and error bars indicate the mean and standard deviation. ****P≤0.0001, ***P≤0.001, ^n.s.^P>0.05; Ordinary one-way ANOVA with Tukey’s method for multiple comparisons. Scale bar = 5μm. Strain imaged was ARF404.

As described, homotypic fusion follows the last round of asymmetric division and precedes the onset of the final (L4/A) molt (Podbilewicz and White, 1994) (Fig. 4A). The seam syncytia go on to make the adult-stage alae during the molt. As shown in Fig. S1, the rectangular seam cells of late L3 larvae were connected to each other and hyp7 by AJs incorporating AJM-1::mCHERRY. UTRNCH::GFP labeled F-actin within cortical meshworks and alongside AJs. The seam cells became more ellipsoidal during the L3/L4 molt and divided shortly thereafter. The anterior daughters fused with hyp7 as the posterior daughters reconnected (Fig. S1). Surprisingly, we found seam syncytia, rather than individual cells, in all nematodes imaged 3-hours after emergence in L4 (Fig. 4B). Capturing the dynamic redistribution of cortical actin during asymmetric division, cytokinesis, migration of the posterior daughters and homotypic fusion of the seam cells validated our key reagents and approach.

Straightaway, the apical surface of seam syncytia narrowed, as AJs demarcating the dorsal and ventral margins drew closer together (Fig. 4B, 3hrs). The apical surface of the seam then widened slowly, as the AJs spread apart over the following 4-5hrs. This interval included much of the L4/adult molt (Fig. 4B, molt). The observed shape changes indicated a pulse of anisotropic constriction on the D-V (transverse) axis followed by gradual relaxation. Indeed, at the apex of constriction 4hrs into the L4 stage, the width of the seam was 1.9μm ±0.3μm (mean and SD), compared to 3.2μm ±0.7μm before constriction (3hrs in L4) and 2.8μm ±0.4μm at 6hrs in L4. As described below, concurrent changes in the ultrastructure of the epidermis detected by transmission electron microscopy (TEM) further substantiated this model (see Fig. 6).

**Figure 5.**
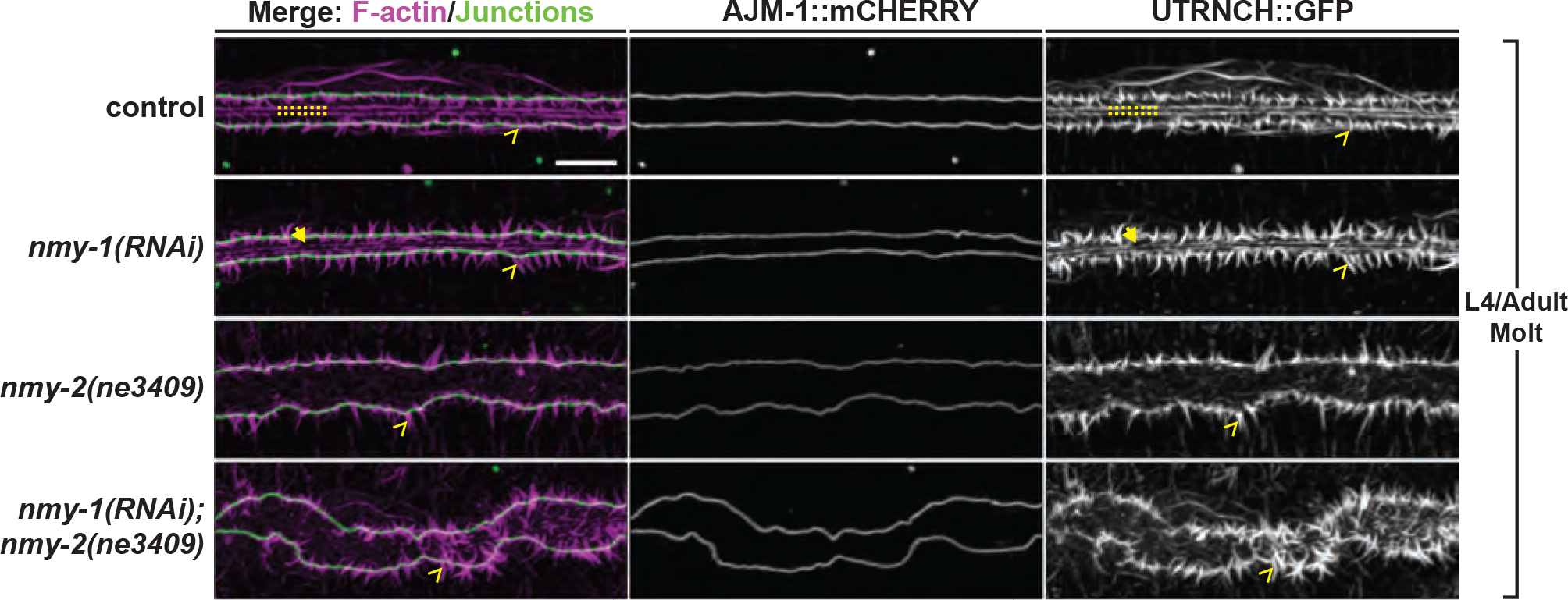
Contractility is required to maintain seam shape. Confocal fluorescence projections show F-actin and AJ signals in worms NMYll knockdowns transiting the L4/Adult molt. Arrowheads point to small breaks in medial-axial AFBs. lmages were acquired using a Zeiss LSM880 with Airyscan processing. Scale bar = 5μm. Strains imaged were ARF404 and ARF410.

**Figure 6.**
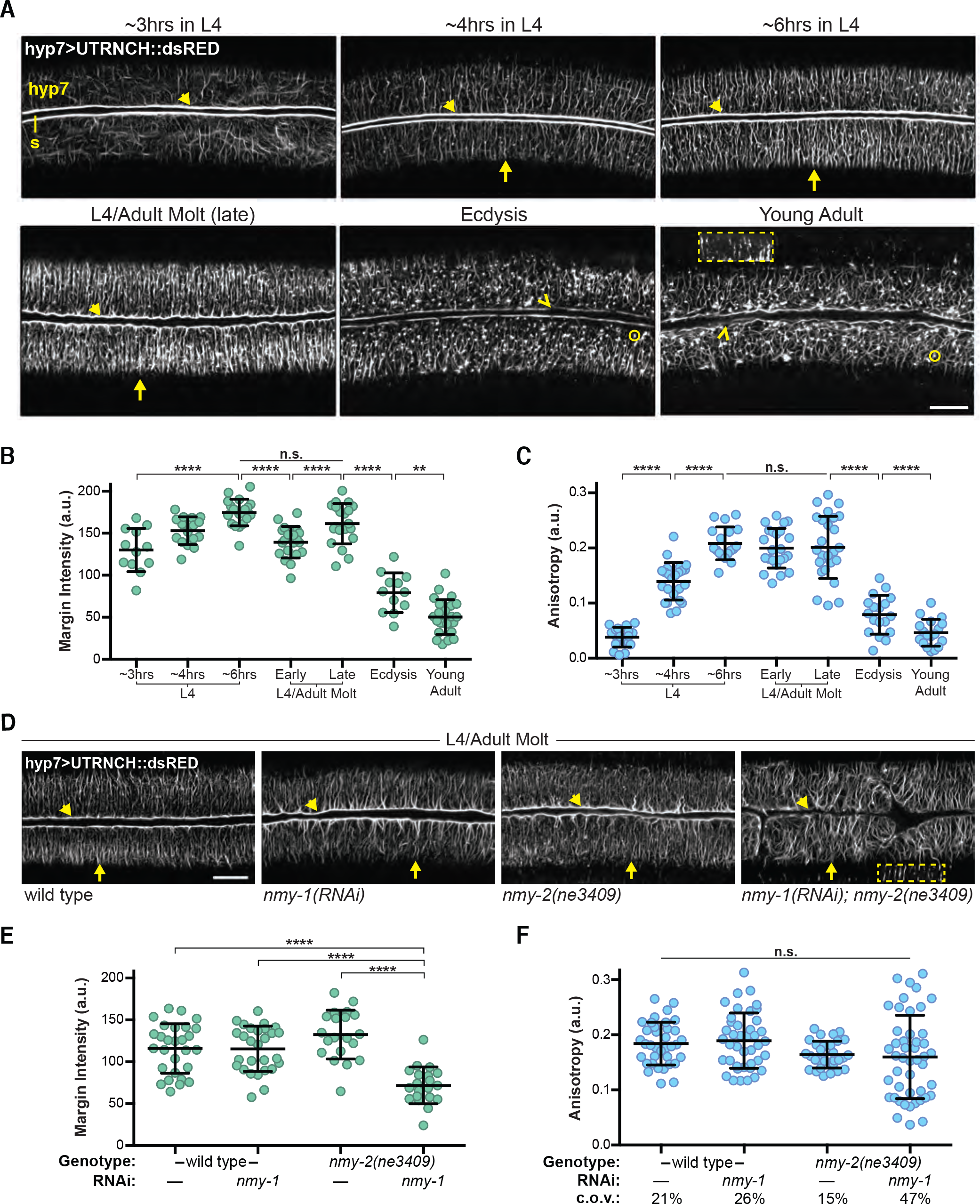
Cortical actin dynamics in lateral regions of hyp7. **A)** Representative confocal fluorescence projections show F-actin labeled by hyp>UTRNCH::dsRED across the larval-to-adult transition. Arrowheads point to F-actin at margins adjoining the syncytial seam; chevrons point to segments with little or no detectable signal. Arrows point to circumferential actin filament bundles (CFBs). Circles mark puncta. Digitally-brightened ROl (2X exposure) shows CFBs in the region of hyp7 overlying body wall muscles. These micrographs were selected from images of over 130 precisely staged specimens. **B)** Quantitation of F-actin signals at the margins of hyp7. Lines and error bars indicate the mean and standard deviation. ****P≤0.0001, **P≤0.01, ^n.s.^P>0.05; Ordinary one-way ANOVA with Tukey’s method for multiple comparisons. **C)** Quantitation of the anisotropy of F-actin signals in laterals region of hyp7. Values for signal intensity and anisotropy represent 4 and 6 ROls per worm, respectively; N=3-7 per stage and condition. **D)** Confocal fluorescence projections show signals from hyp7>UTRNCH::dsRED in molting animals of the indicated genotypes. **E)** Quantitation (as in C) of corresponding F-actin signals at hyp7 margins. **F)** Quantitation of the anisotropy of F-actin signals (as in E). Scale bars = 10μm. Strains imaged were ARF385 and ARF388.

Two longitudinal actin bundles rapidly assembled at the cortex of seam syncytia, co-localizing with AJs along the dorsal and ventral margins (Fig. 4B, arrows). These particular actin bundles persisted as the seam narrowed. The extent to which longitudinal actin bundles overlapped with AJs diminished as the seam widened. Instead, we detected spikes of F-actin orthogonal to the margins (Fig. 4B, chevrons). These spikes increased in size but decreased in number as larvae entered the molt and emerged as young adults. This dramatic reorientation of marginal F-actin relative to AJs suggested a corresponding change in the routing of forces between the seam and hyp7.

Two more actin bundles assembled in the middle of seam syncytia (Fig. 4B, dashes). These medial-axial bundles moved closer together as the seam narrowed, spread apart as the seam widened, and endured the process of molting. Cortical meshworks were detected at all times but relatively difficult to discern in narrow syncytia. At the apex of apical constriction, the pattern of longitudinal actin cables near the surface of seam syncytia bore a striking resemblance to the adult-stage alae, manifest ∼6hrs later. To summarize, we detected four longitudinal actin bundles at 3hrs in L4, prior to seam constriction. At the apex of constriction (4hrs in L4), we detected three longitudinal actin bundles. As the seam widened again, two medial-axial bundles persisted, whereas F-actin spikes appeared at the junctions.

To identify cortical actin structures that most likely generate or route mechanical forces relevant to patterning the alae, we considered that tension *per se* stabilizes actomyosin filaments and related adhesive complexes (Borghi et al., 2012; Hannezo et al., 2015; Wu and Yap, 2013). For this reason, we examined the distribution of AJs and cortical actin in NMYII knockdowns transiting the L4/A molt (Fig. 5). AJs demarcating the dorsal and ventral margins of seam syncytia were tortuous but typically farther apart in *nmy-1(RNAi); nmy-2(ne3409ts)* double mutants than age-matched transgenic nematodes. Distension of the seam syncytia seemed to account for the girth of cuticular mazes found on *nmy-1(RNAi); nmy-2(ts)* adults (Fig. 2A). Further, medial-axial actin bundles and cohesive cortical meshworks were not detected in the seam syncytia of *nmy-1(RNAi); nmy-2(ne3409ts)* double mutants, but were found in control specimens (Fig. 5, dashes). Distended seam syncytia lacking medial-axial actin bundles and cohesive cortical meshworks were likewise observed in *nmy-1(RNAi); nmy-2(ne1409ts)* double mutants. Qualitatively similar but less severe abnormalities were observed in the corresponding single mutants (Fig. 5). In contrast, spikes of F-actin orthogonal to AJs were readily detected in *nmy-1 nmy-2* double and single knockdowns, as well as control specimens. Thus, the assembly and/or persistence of medial actin cables and meshworks near the apical surface of seam syncytia depends on NMYII more than the assembly and/or persistence of peripheral spikes. Taken together, these findings suggest that cortical tension contributes to apical constriction of the seam syncytia, its measured relaxation and/or maintenance of the resulting shape of seam syncytia, which are distinct but likely interdependent processes.

### Tension-dependent recruitment of F-actin to the margins of hyp7

As our initial findings indicated that actomyosin networks in both the seam and hyp7 syncytia contribute to morphogenesis of the alae, we went on to track the distribution of cortical actin in hyp7 across the larval-to-adult transition (Fig. 6, Fig. S2). For this purpose, we constructed a similar but distinct F-actin sensor comprising UTRNCH tagged with dsRED and driven by the hypodermal-specific minimal promoter of the cuticle collage gene *dpy-7* (Johnstone et al., 1992). Longitudinal actin bundles were detected along the lateral margins of hyp7 throughout L4 and the L4/A molt (Fig. 6A & B). The intensity of corresponding signals peaked ∼6hrs after larvae emerged in the L4 stage and plummeted as adults escaped from the larval cuticle. Spikes of F-actin orthogonal to the lateral margins were also detected in molting animals. To evaluate the possibility that actin at the margins of hyp7 regulates changes in seam cell shape, we examined these structures after genetic inactivation of NMYII. Signals from hyp7>UTRNCH::dsRED were barely detectable at the boundary between hyp7 and seam syncytia in *nmy-1(RNAi); nmy-2(ne3409)* double mutants molting from L4 to the adult stage (Fig. 6D). The average fluorescence intensity at this boundary was 72±22 a.u. (N=5) in double mutants, compared to 116±20 a.u. (N=7) in transgenic but otherwise wild type animals (Fig. 6E). Finding that actomyosin contractility maintains actin bundles on the hyp7 side of AJs suggests that these force-transmitting complexes regulate the pulse of apical constriction of the seam syncytia during the larval-to-adult transition.

Cortical meshworks were detected in the lateral (thick) regions of hyp7 syncytia ∼3hrs after larvae emerged in the L4 stage. Parallel arrays of circumferentially-oriented actin filament bundles (CFBs) assembled near the apical surface of hyp7 over the following 3hrs; persisted through the molt; and then abruptly collapsed. These dynamics were substantiated by anisotropy values (a metric for parallel alignment) of UTRNCH::dsRED signals in this region of interest. On average, anisotropy values increased 5-fold during the L4 stage, remained high for most of the L4/A molt, plummeted at ecdysis, and returned to base-line in young adults (Fig. 6C). We further examined the status of CFBs in hyp7 upon inactivation of NMYII. CFBs persisted in virtually all *nmy-1(RNAi); nmy-2(ne3409)* double mutants and corresponding single mutants molting from L4 to adult (Fig. 6D). In one double mutant, neither the seam syncytium nor CFBs were detected, indicating an atypically severe phenotype. Consistent with these observations, no significant differences were found among mean anisotropy values for UTRNCH::dsRED in mutants versus control (Fig. 6F). Thus, neither the assembly or maintenance of CFBs in lateral regions of hyp7 was obviously dependent on actomyosin contractility. Moreover, the increased variability in anisotropy values associated with NMYII knockdown was not mirrored by irregular patterning of the annuli (Fig. 2C). As such, it is unlikely that CFBs directly pattern the annuli or propagate their pattern from L4 to adult-stage cuticles. The kinetics of CFB assembly and disassembly further suggested an alternative function.

### Transient CFBs in lateral hyp7 connect to stable CFBs in hyp7 over muscle

During the above-mentioned study, we noticed that some of the transient CFBs detected in the lateral region of hyp7 across the L4-to-adult transition extended into the quadrant of hyp7 overlying body wall muscle (Fig. 6D and Fig. S2). These previously unrecognized CFBs in hyp7 overlying body wall muscle (mCFBs) were consistently observed in L4, molting animals, and adults (Fig. S2A and B). The mCFBs were situated between *Ce.* hemidesmosomes (*Ce*HDs) (Fig. S2B and C) labeled either by MUP-4::GFP, which is an affiliated apical ECM receptor (Hong et al., 2001) or VAB-10A::GFP, which is the core *Ce*HD spectraplakin (Morrissey et al., 2014). These findings suggest that mCFBs are components of stable attachment complexes distinct from *Ce*HDs.

### Ultrastructure of the epidermis during deposition of the alae

Rapid constriction and slow expansion of seam syncytia regulated by actomyosin networks hinted at reversible folding of the apical surface over the larval-to-adult transition. We therefore used transmission electron microscopy to further characterize related changes in the ultrastructure of the lateral epidermis and overlying matrices.

The TEM micrographs in Figure 7 show transverse sections through the lateral epidermis of early versus mid L4 larvae presented in initial studies of the molting cycle and reproduced with permission (Singh and Sulston, 1978). The apical surface of the seam narrowed midway through L4. The width of the seam decreased by approximately 3.5-fold as indicated by the distance between the two apical-lateral junctions with hyp7. Three discrete protrusions (blebs) were detected on the apical surface. Two blebs were situated above seam-hyp7 junctions, and one near the middle of the seam syncytium. A thin but continuous ECM was detected above the epidermis and beneath the preexisting larval cuticle, covering all three cellular protrusions. This distinctive apical matrix almost certainly corresponds to the molting sheath recognized some 40yrs later (Katz et al., 2015).

**Figure 7.**
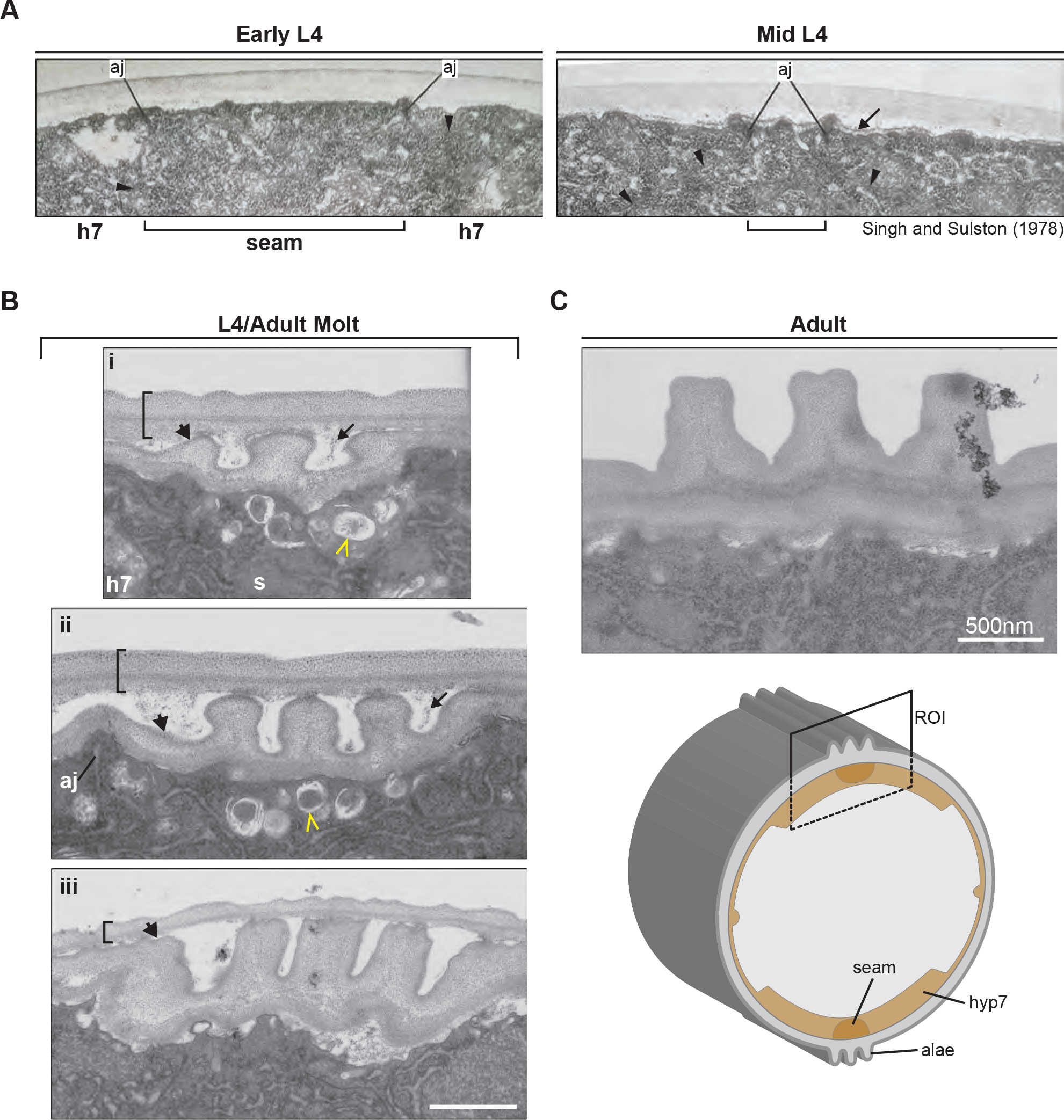
Changes in tissue ultrastructure coupled to morphogenesis of the alae. **A)** TEM micrographs show transverse sections through the seam and hyp7 syncytia of early and mid-L4 wild-type larvae (25,000X). Brackets demarcate the width of the seam surface; aj = apical junctions. Arrowheads outline the seam syncytium. The singular arrow points to the provisional sheath situated above cellular protrusions and beneath the L4 cuticle. Adapted with permission from Singh and Sulston (1978) (Singh and Sulston, 1978). **B)** Comparable TEM micrographs across the L4/A molt show invagination and expansion of the seam cortex during deposition of the alae. Brackets demarcate effete larval cuticles; arrowheads point to emergent adult cuticles; arrows point to presumptive remnants of the sheath. Chevrons point to secretory granules in the seam. **C)** Transverse TEM section shows the alae and lateral epidermis of a young adult at the same magnification. Scale bar = 500nm. The complementary illustration depicts the epidermal syncytia and mature alae of an adult worm from the same perspective.

Inward deformation and gradual expansion of the apical membrane of the seam syncytium followed during the L4-to-adult molt, producing a large pocket in which the alae were deposited (Fig. 7B). As the seam widened, the seam-hyp7 junctions moved farther away from the nascent alae, whose position on the D-V axis did not change appreciably across the molt. Progressive deposition of the alae beneath the L4 cuticle and remnants of the sheath were indicated by the increasing height of the ECM ridges. Secretory granules were present within the seam syncytium, below the growing alae and intervening segments of the apical membrane. Notably, the L4 cuticle thinned as the alae grew, suggesting that the effete cuticle was partially dismantled and possibly recycled. In young adults, the surface of the lateral epidermis followed the contour of the body and the ridges of the tripartite alae were apparent (Fig. 7C).

### Provisional sheaths relay cortical actin dynamics to fixed patterns in apical matrices

Based on the spatial and temporal correlations among the longitudinal actin bundles present in seam syncytia at the apex of constriction (Fig. 4B); transient protrusions from the apical surface of seam syncytia (Fig. 7A); and the appearance of longitudinal bands in the provisional sheaths of animals molting from L4 to the adult stage (Fig. 8), we hypothesized that cortical actin dynamics were mechanically coupled to this short-lived aECM. As an initial test of this hypothesis, we determined the effect of attenuated *actin* RNAi on the pattern of two components of the L4/A sheath; namely, the ZP-domain proteins FBN-1 and NOAH-1. Both *fbn-1* and *noah-1* emerged from a preliminary, RNAi-based screen for knockdowns unable to molt (Frand et al., 2005). Thereafter, we constructed full-length (FL) *fbn-1::mcherry* and *noah-1::sfgfp* translational fusion genes within distinct extrachromosomal arrays (*aaaEx78* and *aaaEx167*). These arrays rescued the lethality caused by respective null alleles of *fbn-1* or *noah-1*, confirming the production of functional fusion proteins. As previously described and shown in Fig. 8A, circumferential bands of FL-FBN-1::mCHERRY corseted the body while larvae underwent molts and while embryos elongated. Longitudinal bands of FL-FBN-1::mCHERRY were also detected above the seam syncytia during the L4/A molt. The pattern of these bands presaged the forthcoming alae (Katz et al., 2015). Longitudinal and circumferential bands of NOAH-1::sfGFP were also detected above the seam syncytia during the L4/Adult molt (Fig. 8B). In this case, the fluorescent signal localized to four distinct stripes aligned with light-refractive regions of the nascent cuticle visible by DIC; these stripes likely correspond to temporary valleys flanking the emergent alae.

**Figure 8.**
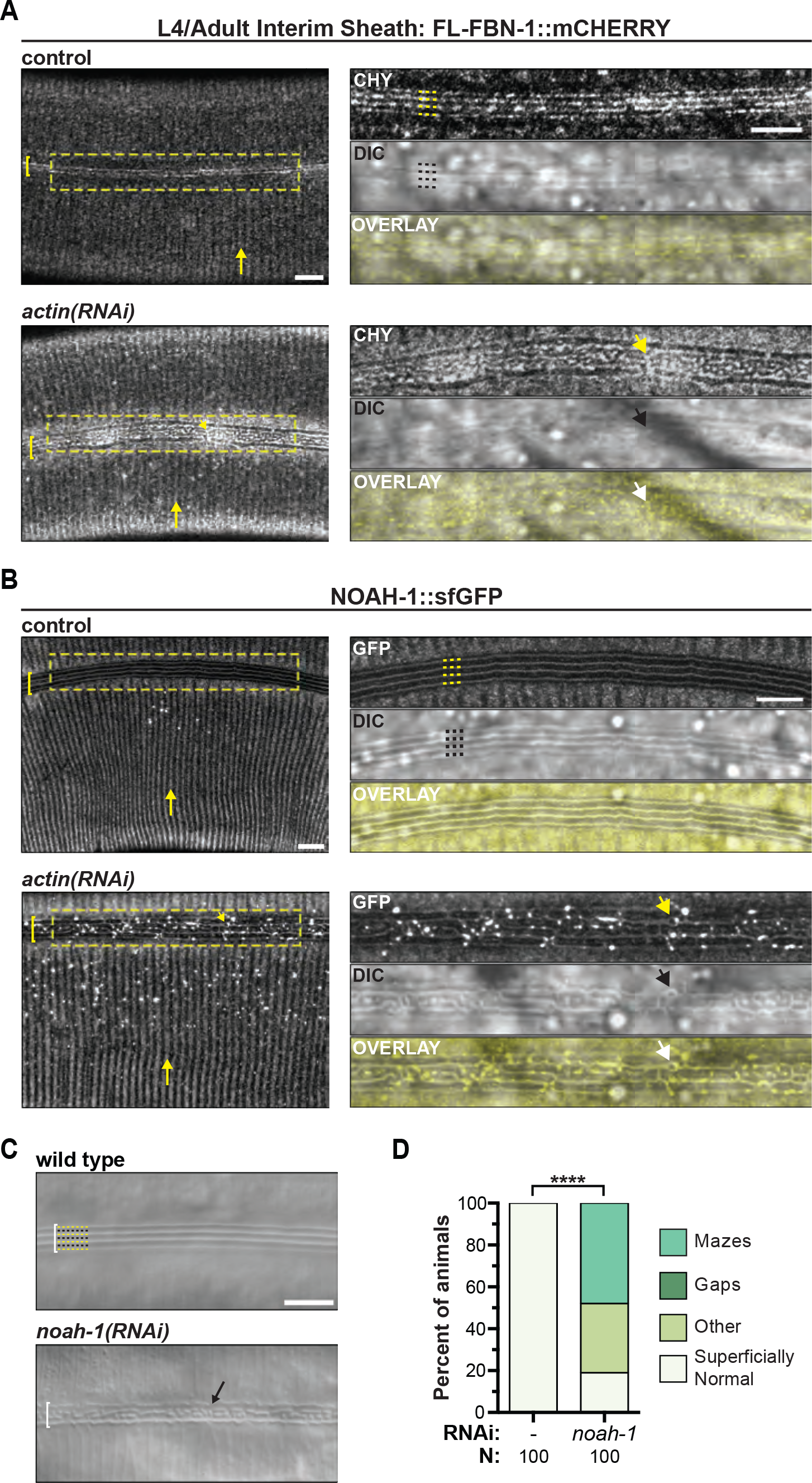
Actin is required to pattern sheath components, which then pattern the alae. **A)** Representative confocal fluorescence projections show FL-FBN-1::mCHERRY detected in the sheaths of control and attenuated *actin(RNAi)* animals molting from L4 to the adult stage. Vertical arrows mark fluorescent annular bands within the sheath. Brackets demarcate the underlying seam syncytia. Dashed rectangles indicate ROIs digitally enlarged in adjacent panels. In control, ROI corresponds to a maximum intensity projection omitting surface slices. Yellow dashes label fluorescent longitudinal cables detected on the lateral surface of the control specimen; black dashes label the same sites on the corresponding DIC micrograph. Arrowheads point to fluorescent maze-like fibrils detected on the lateral surface of the attenuated *actin(RNAi)* specimen and similar substructures on the emergent cuticle. Fluorescent signals were false colored yellow to make the overlays. Images were acquired using a Zeiss LSM880 with Airyscan processing. **B)** As described, NOAH-1::GFP primarily localizes to the sheath. These representative confocal projections show residual signals detected on the surface of newly emerged control and attenuated *actin(RNAi)* adults, essentially as in A. **C)** Representative DIC micrographs show the lateral surfaces of young adults. Black and yellow dots label longitudinal ridges (alae) and adjacent valleys on the control specimen. Arrow points to a maze on the *noah-1(RNAi)* animal. **D)** Quantitation of alae defects. As previously described, values are the weighted average of 2 independent trials. ****P<0.0001; Fisher’s exact test. Scale bars = 5μm. Strains used were ARF379, ARF415, and N2.

Following attenuated RNAi of *actin*, FL-FBN-1::mCHERRY localized to labyrinthine deformities, rather than longitudinal bands, overlying the lateral surfaces of emergent adult-stage cuticles (Fig. 8A). The abnormal pattern of FL-FBN-1::mCHERRY within the sheath matched the abnormal pattern at the surface of emergent cuticles detected by DIC. Attenuated RNAi of *actin* also led to the deposition of NOAH-1::sfGFP mazes perfectly aligned with misshapen valleys in the emergent cuticle above the seam syncytia (Fig. 8B). In contrast, the annular patterns of FL-FBN-1::mCHERRY and NOAH-1::sfGFP were superficially normal in *actin(RNAi)* animals, consistent with our initial findings that both epidermis-specific and systemic but attenuated RNAi of *actin* impaired *de novo* patterning of the alae, rather than propagation of the pattern of annuli.

The above-mentioned hypothesis further predicts that knocking-down certain sheath components will lead to deformities in apical ECMs. We therefore examined the lateral surfaces of young adults following attenuated but systemic RNAi of *noah-1*. Mazes and minor deformities in the alae were detected in 81% (N=100) of *noah-1(RNAi)* adults, but were not detected in same-day controls, consistent with our working model that provisional sheaths propagate cortical actin dynamics underlying transient changes in epithelial cell shape to sculpt more durable apical matrices.

Having conceived this working model and validated its key predictions in the context of the larval-to-adult transition, we asked whether a similar, if not identical, morphogenetic mechanism gives rise to the architecturally distinct alae of dauers. To address this question, we combined the conditional *daf-2(e1370)* allele, which confers a dauer-constitutive (Daf-c) phenotype at restrictive temperature, with our seam>UTRNCH::GFP sensor for F-actin and our FL-FBN-1::mCHERRY marker for the provisional sheaths of molting worms. We then imaged UTRNCH::GFP in live nematodes molting from L2d to dauer, a process that involves extreme radial constriction of the body. Early in the molting process, dense cortical meshworks including both longitudinal and transverse actin filaments and/or bundles were detected in rather wide, seam cells (Fig. S3A). Some of these structures appeared supracellular, crossing cell boundaries, or were aligned in adjacent cells. Later in the process, we detected spikes of actin orthogonal to junctions between seam cells and hyp7 in the very thin seam cells of fully constricted animals. We also imaged FL-FBN-1::mCHERRY across the L2d to dauer molt. The pattern of FL-FBN-1::mCHERRY in the corresponding sheath included five longitudinal stripes within a wide lateral band (Fig. S3B). While this pattern was markedly different from the sheaths of animals molting from L4 to the adult stage, the pattern of FBN-1 stripes yet again presaged the shape of the forthcoming dauer-specific alae, which have four ridges that span most of the lateral surface of the cuticle (Fig. S3C). In addition, the lack of detectable FL-FBN-1::mCHERRY signal on the surface of dauers was consistent with the temporary nature of the sheath, which was partly consumed and partly shed with the L2 cuticle. These observations are entirely consistent with temporal reiteration of a common morphogenetic mechanism.

## DISCUSSION

Figure 9 provides cohesive model for morphogenesis of the lateral epidermal syncytia and cuticle of adult-stage *C. elegans*, based on key findings from descriptive and mechanistic studies. Therein, rearrangements in cortical actin networks effect transitory changes in cell shape and tissue architecture, which are relayed through the provisional sheath to pattern permanent ridges on the cuticle. Midway through the 4^th^ larval stage, the seam syncytium narrows rapidly on the dorsal-ventral axis (Fig. 9A 1-2). Forces generated by actomyosin filaments and propagated through adherens junctions evidently drive this change in shape, as knockdowns of epidermal actin, NMYII, and HMP-1 all give rise to distended seam syncytia and/or mazes. Longitudinal actin bundles assemble on both sides of seam-hyp7 junctions during this phase; as described below, these parallel bundles may act cooperatively to drive AC. Next, the seam syncytium widens slowly, suggesting the gradual release of cortical tension. Such relaxation is atypical among morphogenetic processes dependent on AC, as molecular ratchets often fix the shapes of constricted cells (Mason et al., 2013). Spikes of F-actin orthogonal to seam-hyp7 margins arose during the relaxation phase. Related adhesive complexes may allow the seam to push back against hyp7 (Fig. 9A 2-3) and so resemble dynamic endothelial junctions (Cao et al., 2017; Lampugnani, 2010).

**Figure 9.**
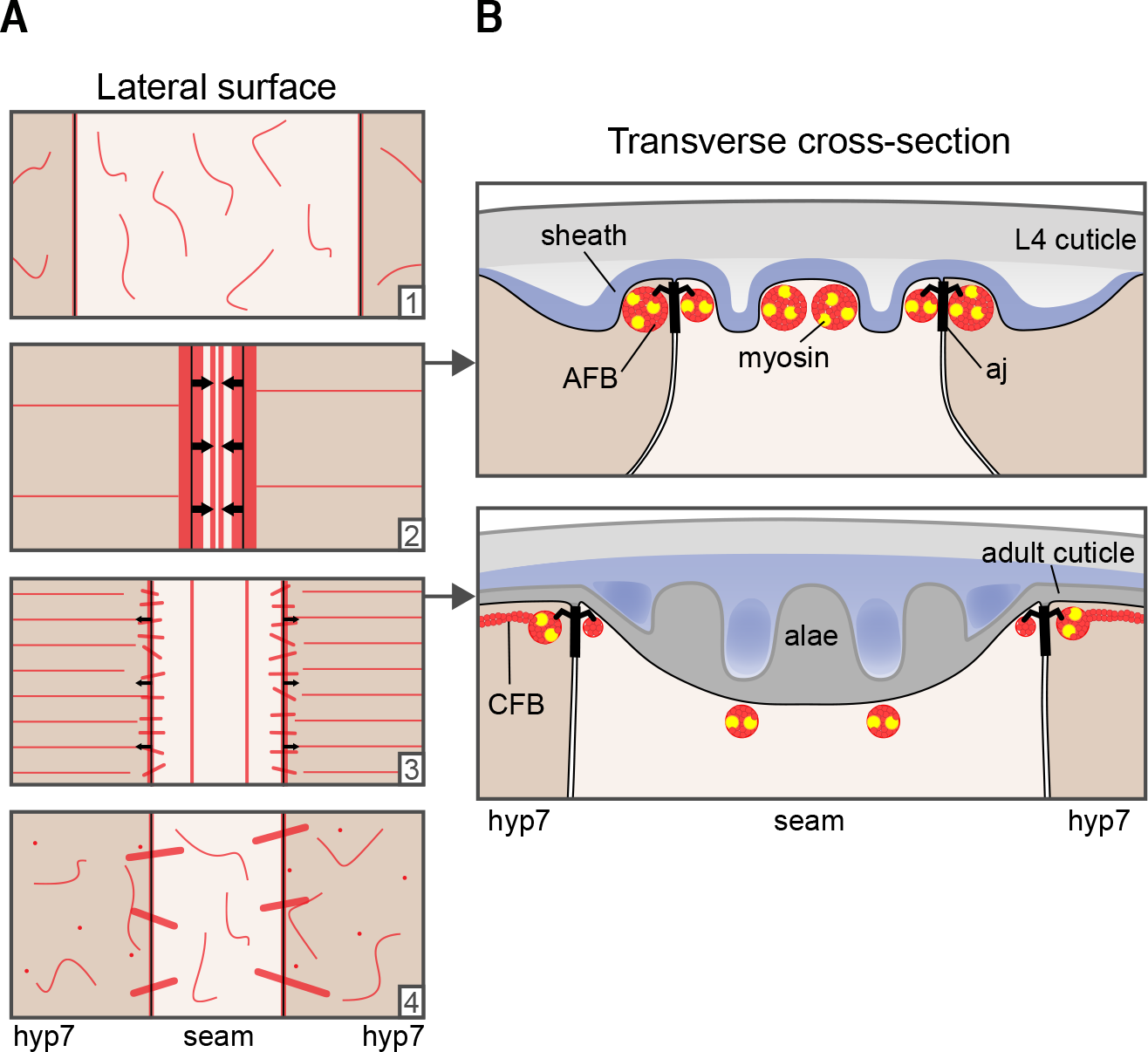
Synopsis of key findings and model for morphogenesis of the adult-stage alae. **A)** Graphical representation of the lateral surface of the seam and hyp7 syncytia across the larval-to-adult transition. Cortical actin networks are depicted in red. Inward facing arrows (2) represent apical constriction of the seam syncytium driven by net compressive forces. Outward facing arrows (3) represent gradual expansion of the seam controlled by the dynamic exchange of mechanical forces among epidermal syncytia and the molting sheath. **B)** Graphical representation of a transverse cross-section through the lateral epidermis and overlying matrices at the apex of constriction (A2) and subsequent relaxation (A4) of the seam cortex. The related model for patterning the alae is fully described in the discussion.

The forces that drive constriction of the seam syncytium also generate three cellular protrusions via outward folding of the apical membrane: one above the dorsal junction with hyp7, one in the middle of the seam syncytium, and one above the ventral junction (Fig. 9B, right). These protrusions effectively mark the sites of forthcoming ridges in the adult cuticle. The outer blebs spatially and temporally correspond with longitudinal actin bundles that co-localize with AJs. The middle bleb probably forms above the medial-axial actin bundles, which persist well into the molt. The blebs are another noteworthy aspect of this system, given that AC typically leads to inward cellular deformations and tissue invagination (Heer and Martin, 2017), with few exceptions such as budding of the lungs (Kim et al., 2013). However, F-actin has been detected in cellular blebs during the formation of taenidial folds in the *Drosophila* trachea, as the chitinous aECM is deposited {Ozturk-Colak, 2016 #121}.

As described, all three protrusions appear beneath the molting sheath, which is a temporary extracorporeal enclosure composed of several ZP-domain proteins. Although the blebs are ephemeral, the pattern of the lateral sheath mimics the cellular protrusions and the forthcoming alae. In this context, the sheath relays forces from passing actin networks to shape upcoming substructures in durable cuticles. This could occur through transient coupling of the actin cytoskeleton to specific sheath components by a currently unknown transmembrane receptor. Consistent with this model, attenuated knockdown of *actin* abrogates the pattern of longitudinal cables within the sheath visualized by FBN-1 and NOAH-1 fusion proteins. In turn, knockdown of *noah-1* gives rise to mazes, rather than continuous ridges (alae), on the cuticles of young adults. This last finding identifies the sheath as a key intermediate in morphogenesis of the alae, a process that begins with AC and overcomes delays linked to systemic tissue remodeling.

Three prospective properties would enable the sheath to transmit the pattern of cellular blebs to the ridges of the alae: cell-aECM adhesion, mechanical resilience and space-filling. In theory, adhesion between the cellular protrusions and the sheath could preserve their form while the seam begins to relax. Potential receptors include MUP-4 and MUA-3, which are nematode-specific apical ECM receptors associated with CeHDs (Bercher et al., 2001; Hong et al., 2001) and the two dimeric integrins expressed by *C. elegans* (Cox and Hardin, 2004), which might interact with the canonical integrin-binding motifs present in FBN-1. In this system, mechanical forces generated by the actomyosin cytoskeleton in the underlying epidermis are routed through the sheath. Related tension-bearing capacity could be provided by organization of ZP-domain meshwork or resilience of specific components therein. Lastly, the sheath occupies space, filling the pocket created by inward deformation of the seam membrane, consistent with O-linked glycosylation of ZP-domain proteins (Gupta et al., 1999). The sheath is dismantled as the alae are progressively secreted, and this may account for accretion of related fluorescent signals in the valleys between and alongside ridges of the alae approaching ecdysis.

While we do not yet know the extent to which longitudinal actomyosin filament bundles (AFBs) within the curved cortex of seam syncytia drive constriction on the transverse (D-V) axis, rather than cortical meshworks; we recognize two applicable mechanisms. First, cross-linking of longitudinal AFBs to one another and/or cortical meshworks would allow for transverse propagation of contractile forces. Prospective cross-linkers include alpha-actinin and formin, which regulate actin dynamics in other instances of AC (Mason et al., 2013). Second, the syncytial seam can be modeled as a sealed cylinder with internal, pressurized fluids. As such, the predicted effect of axial stress is buckling (continuous blebs) on the A-P axis (Nelson, 2016), and the production of circumferential and transverse force vectors (Hayashi et al., 2018; Shih et al., 2017), which may promote anisotropic constriction or bear tension to resist excessive expansion. Modeling the seam syncytium in this way further suggests that cortical flows may determine the initial position of the medial-axial AFBs, which assemble at predicted regions of high tension.

Observing breaks in the medial-axial AFBs and later the middle ridges of *nmy-1(RNAi)* adult-stage alae further implicates this set of AFBs. During the L4-to-adult molt, these AFBs may additionally resist excessive deformation of the apical membrane of the seam under the weight of the growing alae. Manipulations that prevent the appropriate routing of forces would give rise to uneven localized tension at the apical membrane, disordered blebs and subsequent alae (mazes). In addition, insufficient resistance to axial stress would result in transverse breaks. Similar specialized F-actin structures in the seam are not detected in immature seam cells or during prior larval-to-larval molts.

CFBs beneath the apical membrane of hyp7 assemble while the seam widens and persist through most of the molt. The CFBs were not sufficient to produce patterned alae, as CFBs were detected in NMYII knockdowns associated with mazes. However, the formal possibility remains that transient CFBs affect relaxation of the seam.

Knocking down core components of actomyosin networks had no appreciable effect on the fundamental pattern of the annuli, suggesting that the CFBs in hyp7 are largely dispensable for propagating the pattern of annuli from larval to adult-stage cuticles. In addition, variability in the pattern of CFBs was not mirrored by abnormal patterning of the circumferential annuli. Instead, transient CFBs detected in the lateral regions of hyp7 during the L4 to adult transition are temporary elaborations of actin cables present in dorsal-lateral and ventral-lateral quadrants of hyp7 and situated between the apical hemidesmosome-like plaques of fibrous organelles at all times. In theory, these actin cables may be coupled to the apical membrane and thereby components of the musculoskeletal system. One possibility is that these CFBs interact with the actin-binding spectraplakin VAB-10B, which is similarly situated between the *Ce*HDs of late embryos and larvae (Bosher et al., 2003).

Three aspects of alae morphogenesis bear similarity to embryonic elongation: large-scale epidermal actin cytoskeleton rearrangements; the involvement of a temporary sheath capable of propagating mechanical forces; and compaction of the body, albeit to very different extents. Elongation is driven by forces generated by actomyosin contractility in the seam and distributed through CFBs in the hypodermis as well as the sheath (Vuong-Brender et al., 2017a; Vuong-Brender et al., 2017b). The CFBs present in the hypodermis of embryos are morphologically similar to those we describe at the L4-to-adult transition. However, our results suggest that forces generated by the longitudinal AFBs at the junctions between the seam and hyp7 route stress through the sheath and not CFBs. The larval sheath also appears to be similar in composition and function to its embryonic counterpart. For example, FBN-1 and NOAH-1 are components of both (Kelley et al., 2015; Vuong-Brender et al., 2017b). However, the role of the embryonic sheath in L1 alae morphogenesis has not been determined. Compaction is most striking in elongation, when the embryo transforms from a ball of cells into a long, thin, tapered worm (Priess and Hirsh, 1986). In contrast, compaction during the molts has been reported (Uppaluri and Brangwynne, 2015), but seems temporary to accommodate enlargement of the cuticle. The transition from L2d to dauer may present an exception, as radial constriction requires ZP-domain proteins called cuticulins, some of which are found in dauer alae while others may localize to the L2d/dauer sheath (Sapio et al., 2005).

These findings uncover a novel but likely conserved morphogenetic mechanism that links mechanical networks driving apical constriction with matrix dynamics. Given that the key intracellular and extracellular molecules are conserved between nematode and mammals, this new system is likely relevant to mammalian physiology and pathology.

## MATERIALS AND METHODS

### Strains and Molecular Biology

*C. elegans* strains used in this study are listed in Table S1. Strains were maintained under standard conditions and cultivated at 25°C unless otherwise specified (Brenner, 1974). For synchronization, gravid worms were bleached to isolate eggs, hatchlings arrested in starvation-induced L1 diapause, and released from diapause by plating on NGM seeded with *E. coli* 0P50-1. Selected behaviors were scored to bin worms prior to mounting them on slides: locomotion and feeding indicated non-molting worms; quiescence indicated molting worms (Cassada and Russell, 1975); and idiosyncratic movements used to escape larval cuticles indicated ecdysis (Singh and Sulston, 1978). The precise stages of L4 and adult worms were further determined based on the shape of the gonad, vulva and lateral epidermis (Hubbard and Greenstein, 2005; Mok et al., 2015; Sulston and Horvitz, 1977). Cuticle caps over the mouth were indicative of worms undergoing molts (Monsalve et al., 2011). Strains with conditional alleles were propagated at permissive temperature (15°C) and cultivated at restrictive temperature (25°C) following release from starvation-induced L1 diapause. Bacterial-mediated RNA-interference (RNAi) was performed as described (Fraser et al., 2000; Kamath and Ahringer, 2003; Timmons et al., 2001), except that NGM (nematode growth medium) plates were supplemented with 8mM, rather than 1mM, isopropyl β-D-1-thiogalactopyranoside (IPTG). For attenuated RNAi treatments, animals were washed off from control plates 14hrs after release from L1 diapause with 14ml M9, rotated for 30 minutes in M9 to remove residual gut bacteria, and then transferred to experimental RNAi plates. As a control, worms were fed the same *E. coli* HT115(DE3) transformed with pPD129.36. Upon induction by IPTG, such bacteria produce short dsRNA molecules that does not match any annotated gene of *C. elegans*.

Table S2 describes the oligonucleotides used in this study. Phusion High Fidelity Polymerase (NEB) was used to amplify DNA for sequencing and cloning. Taq Polymerase (NEB) was used for PCR-based genotyping. Gibson assembly (NEB) and standard cloning reactions were used to construct fusion genes and corresponding plasmids. To create the *seam*>*rde-1*+ *elt-5p::rde-1::sl2-mcherry::unc-54* 3’-UTR fusion gene housed in pSK08, the promoter of *elt-5*, which corresponds to nucleotides 1910072-1913471 of chromosome IV (GenBank: NC_003282); the coding region of *rde-1*, which corresponds to nucleotides 9988043-9991614 of chromosome V (GenBank: NC_003283); coding sequence for *mCherry* (GenBank: KT175701), *sl2* (GenBank: LK928133); and the *unc-54* 3’-UTR cassette from pPD95.75 were combined. To construct the *hyp-7*>*rde-1*+ *dpy-7p::rde-1::sl2::nls-gfp::unc-54* 3’-UTR fusion gene housed in pSK38, the minimal promoter of *dpy-7*, which corresponds to nucleotides 7537789-7537869 and 7537914-7538219 of chromosome X (GenBank: NC_003284); the coding region of *rde-1*; *SL2*; and the *nls-gfp::unc-54* 3’-UTR cassette from pPD95.73 were united. To construct the *dpy-7p::utrnch::dsred::unc-54* 3’-UTR fusion gene housed in pSK26, the promoter of *dpy-7* (as above); the coding sequence for the first CH domain (residues 1-261) of human Utrophin (GenBank: LX69086); the coding sequence for *dsRed* (GenBank: HQ418395); the *unc-54* 3’-UTR cassette from pPD95.81; and the pUC57 backbone were combined. To construct the *elt-5p::utrnch::gfp::unc-54* 3’-UTR fusion gene housed in pSK34, the promoter of *elt-5*; the sequence encoding UTRNCH; and the *gfp::unc-54* 3’-UTR cassette and backbone from pPD95.81 were united. All variants of plasmids pPD95 were gifts from Andy Fire.

Towards the production of a full-length *fbn-1::mCherry* fusion gene, genomic DNA spanning 3.6kb of upstream sequence and exons 1-14 of *fbn-1k* (Genbank: JQ990128); cDNA spanning exons 14-22 isolated from pMH281 (Maxwell Heiman, Harvard Medical School); and genomic DNA spanning exons 21-25 and the 3’-UTR of *fbn-1k* were combined in plasmid pSK27. The latter fragment was modified by insertion of an in-frame *NotI-mCherry-NotI* cassette between the codons for H2418 and V2419 of *fbn-1k. mCherry* was isolated from KP1272 (Joshua Kaplan, HMS). A 7.4 kb region of genomic DNA spanning the entire promoter and exons 1-2 of *fbn-1* was amplified separately. pSK27 and the purified PCR product were co-injected at an equimolar ratio, allowing for homologous recombination *in vivo* (see below). The genomic DNA present in the full length (fl) *fbn-1::mCherry* fusion gene corresponds to nucleotides 7619229-7641053 of chromosome III (GenBank: LK928224). To construct the *noah-1::sfGFP::noah-1* translational fusion gene housed in pCM05, regulatory and coding regions of *noah-1* were amplified from genomic DNA (nucleotides 5874389-5883950 of chromosome I, GenBank: LK927608) and cloned into a NotI-filled derivate of pCR-Blunt II-TOPO (Invitrogen). A *NotI-sfgfp-NotI* cassette was inserted in-frame between the codons for P624 and V625 of *noah-1a* (Genbank: NM_170870). The corresponding NotI site was created using a Q5 mutagenesis kit (Invitrogen). Superfolder (sf) GFP was isolated from pCW11 (Max Heiman, Harvard University).

All extrachromosomal arrays were generated by microinjection of young adults with mixtures containing a total of 100ng/¼l DNA. To generate *aaaEx37*, pSK08 (5ng/μl); *ttx-3::gfp* (40ng/μl); and pRS316 (55ng/μl) were co-injected into JK537 *rde-1(ne219)*. To generate *aaaEx162*, pSK38 (5ng/μl); *ttx-3::dsred* (40ng/μl); and pRS316 were co-injected into JK537. To generate *aaaEx108*, pSK26 (0.5ng/μl); *ttx-3::gfp*; and pRS316 were co-injected into N2. To generate *aaaEx117*, pSK34 (5ng/μl); *ttx-3::gfp*; and pRS316 were co-injected into N2. Optimal plasmid concentrations used to generate tandem *utrnch* arrays were empirically determined by titration. UTRNCH signals were readily detected in the resulting transgenic animals, while phenotypes associated with high levels of UTRN were not observed. To generate *aaaEx78[fl-fbn-1::mCherry::fbn-1]*, pSK27 (2.5 ng/μl); the above-mentioned PCR product (1.15 ng/μl); *ttx-3p::gfp*; and pRS316 were co-injected into N2. To generate *aaaEx167*, pCM05 (1ng/μl); *ttx-3::dsred*; and pRS316 were co-injected into ARF379. Resulting transgenic lines were out-crossed to N2 to remove *aaaIs12*. Extrachromosomal arrays were integrated into the genome by UV irradiation at 450 kJ using an FB-UVXL-1000 (Fisher Scientific). Strains with newly-integrated arrays were back-crossed to JK537 or N2 4 to 6 times prior to further analyses.

To knockdown *actin* by bacterial-mediated RNAi, we used the clone for *act-2* present in the Ahringer library (Kamath and Ahringer, 2003). Towards knocking-down *nmy-1* (Genbank: LK927643), 1121bp of genomic DNA from exon 10 was cloned into pPD129.36, the standard expression vector for dsRNAs. For *zoo-1* (GenBank: NM_001026515), cDNA spanning exons 1-7 was cloned into pPD129.36, as previously described (Lockwood et al., 2008). For *noah-1* (GenBank: LK927608), 1024bp from exon 6 was cloned into pPD129.36. Each of the resulting plasmids (pSK43, pSK44 and pCM13) was verified by Sanger sequencing and used to transform *E. coli* strain HT115(DE3).

### DiI Staining of Cuticles

DiI staining to visualize cuticle structures was performed basically as described (Schultz and Gumienny, 2012), except that glass pipettes were used to transfer samples in lieu of Triton X-100. Briefly, approximately 600 adult worms were incubated in 400μl of 30 μg/mL DiI (Sigma) in M9 for 3 hours, shaking at 350rpm. Worms were then washed 1X in M9 buffer, re-suspended in 100μl of M9, and dispensed to a 6cm NGM plate seeded with *E. coli* OP50-1. To remove excess unbound dye, worms were allowed to crawl on the plate for 30min prior to imaging.

### Microscopy and Image Analyses

Worms were anesthetized with sodium azide (2.5%) in M9 buffer and mounted on 2% agarose pads. A Zeiss Axioplan microscope with an attached Hamamatsu Orca ER CCD camera was used for compound microscopy. Images were acquired and analyzed using the software package Volocity 6.3 (PerkinElmer). A Zeiss LSM 5 PASCAL microscope controlled by ZEN 9.0 software was typically used for confocal microscopy. A Zeiss LSM880 laser scanning confocal microscope was used as specified. Measurements were made using Volocity 6.3 (PerkinElmer), ImageJ (Version 1.48v, NIH), and Fiji (Schindelin et al., 2012). To determine furrow spacing from images of Dil-stained cuticles, we used the Find Peaks BAR script in Fiji. This tool identified local minima from linescan plots, also made in Fiji. Three to four linescans were made per image. Seam width was measured in Fiji as the distance between AJM-1::mCHERRY-marked junctions. Six measurements were made per image. Volocity was used to measure fluorescence intensity in manually-selected, similarly-sized regions along the margins of hyp7. Four ROIs were assayed per worm. The ImageJ plugin FibrilTool was used to measure anisotropy (Boudaoud et al., 2014). For each worm assayed, 6 values were obtained by subdividing the lateral region of hyp7 into 3 dorsal and 3 ventral ROIs, each approximately 400μm^2^.

For transmission electron microscopy (TEM), synchronized wild-type (N2) animals were collected, washed once in 8% ethanol and M9, and washed 3 times in PBS over a period of 30 mins. Specimens were suspended in 2.5% glutaraldehyde, 1% paraformaldehyde and 0.1M sucrose in PBS; incubated for 2 hours on ice; and incubated for 16 hours at 4°C. Samples were washed, post-fixed in 1% OsO4, and dehydrated by serial immersion in graded ethanol solutions. Samples were then passed through propylene oxide, embedded in serial steps using mixtures of propylene oxide and Epon 812, and cured at 60°C for 48 hrs. An RMC MTX ultramicrotome was used to cut 60 nm-thick sections, which were stained with uranyl acetate and lead citrate. Sections were observed using a 100CX JEOL electron microscope.

### Statistical Analyses and Image Presentation

GraphPad Prism 6 and Microsoft Excel 15.21 were used for statistical analyses. To perform statistical analyses in Figures 2B, 3B, and 8D, the mazes, gaps, and other phenotypical categories were combined so that and outcomes were classified as abnormal versus superficially normal. All micrographs were prepared for publication using Adobe Photoshop v13.0 and Adobe Illustrator v16.0.

## ACKNOWLEDGEMENTS

Some strains used in this study were provided by the *Caenorhabditis* Genetics Center, which is funded by the NIH Office of Research Infrastructure Programs (P40 OD010440). We are grateful to Margot Quinlan, Alvaro Sagasti, and Larry Zipursky for helpful discussions. We thank Dominic Williams for valuable technical assistance. We also thank Margot Quinlan, John Kim, Eric Miska, and David Sherwood for sharing reagents. A Ruth L. Kirschstein National Research Service Award (GM007185 to SK), UCLA Dissertation Year Fellowship (SK), and a National Science Foundation Award (IOS 1258218 to ARF) supported this research.

## SUPPLEMENTAL MATERIALS

**Supplemental Figure 1. Cortical actin dynamics in seam cells prior to homotypic fusion.** Representative confocal fluorescence projections (lateral view) show signals from seam>UTRNCH::GFP and the AJ marker AJM-1::mCHERRY detected across the L3-to-L4 transition. UTRNCH::GFP is false colored magenta; AJM-1::mCHERRY, green in merged images. Bracket demarcates the space between two posterior daughter cells previously occupied by an anterior daughter cell that fused with hyp7, and bridged by UTRNCH::GFP signal. Arrows points to concentrated UTRNCH::GFP signal at the junction between daughter cells. s=seam; a=anterior and p=posterior daughter cell. Strain imaged was ARF404. Scale bar = 5μm.

**Supplemental Figure 2. CFBs in thin regions of hyp7 interdigitate with hemidesmosomes and some extend over the lateral epidermis while animals molt. A)** Confocal fluorescence projections show signals from hyp7>UTRNCH::dsRED (magenta in merge) and MUP-4::GFP 6hrs into the L4 stage. MUP-4 is a nematode-specific ECM receptor associated with apical CeHDs and corresponding fibrous organelles. The dashed line distinguishes the thick, lateral region of hyp7 overlying the pseudocoelom (lh7) and the thin region overlying body wall muscles (mh7). Arrows points to one example of a continuous CFB that spans both regions and contacts the seam/hyp7 margin. s= seam. **B)** Merged projections of the same 2 signals show intercalated stripes of F-actin and MUP-4 across the L4-to-adult transition. **C)** Confocal fluorescence micrographs show F-actin labeled by UTRNCH::dsRED and Ce.HDs marked by VAB-10A::GFP detected at the surface of mh7 in a young adult. The corresponding linescan shows nonoverlapping peaks. Images were acquired using a Zeiss LSM 880 with linear unmixing of dsRED and GFP signals. Strains imaged were ARF412 and ARF407. Scale bars = 5μm.

**Supplemental Figure 3. Cortical actin and apical ECM dynamics across the L2d-to-dauer transition. A)** Confocal fluorescence projections show F-actin labeled by UTRNCH::GFP in the non-fused seam cells of *daf-2(e1370)* larvae. Dashed lines and arrowheads label longitudinal and transverse actin bundles, respectively, detected early in the L2d to dauer molt. Chevrons point to spikes of F-actin orthogonal to the margins of very narrow cells. **B)** Confocal fluorescence projection shows FL-FBN-1::mCHERRY detected on the surface of a molting *daf-2(e1370)* animal. Yellow dashed lines label distinct longitudinal stripes of FL-FBN-1::mCHERRY, numbered pairwise at right, reflecting bilateral symmetry. Bracket demarcates the presumed underlying seam width prior to constriction. Arrow points to a circumferential band of FL-FBN-1::mCHERRY. (Bottom) Trace of dauer alae in published transverse TEM micrographs {Cassada, 1975 #100} shows characteristic ridges and valleys. **C)**For comparison, this overexposed fluorescence micrograph shows that signals from FL-FBN-1::mCHERRY were not detected on the surface of a dauer larva. The bracket on the corresponding DIC micrograph demarcates the dauer alae. Strains imaged were ARF416, ARF417, and ARF321. Scale bars = 5μm.

**Supplemental Table 1:** *C. elegans* strains used in this study.

**Supplemental Table 2:** Oligonucleotides used in this study.

